# Behavioral and neural fusion of expectation with sensation

**DOI:** 10.1101/2020.07.03.187146

**Authors:** Matthew F. Panichello, Nicholas B. Turk-Browne

## Abstract

Humans perceive expected stimuli faster and more accurately. However, the mechanism behind the integration of expectations with sensory information during perception remains unclear. We investigated the hypothesis that such integration depends on ‘fusion’ — the weighted averaging of different cues informative about stimulus identity. We first trained participants to map a range of tones onto faces spanning a male-female continuum via associative learning. These two features served as expectation and sensory cues to sex, respectively. We then tested specific predictions about the consequences of fusion by manipulating the congruence of these cues in psychophysical and fMRI experiments. Behavioral judgments and patterns of neural activity in auditory association regions revealed fusion of sensory and expectation cues, providing evidence for a precise computational account of how expectations influence perception.

## Introduction

Experience and learning guide perception, allowing for fast and accurate processing of sensory input that is noisy and ambiguous (Hutchinson & Turk-Browne, 2012; Oliva & Torralba, 2007; Panichello et al., 2013). As a result, it has long been suggested that perception may be best understood as a form of probabilistic inferences about the outside world, rather than a veridical representation of sensory inputs (Gregory, 1980; Von Helmholtz, 1866). However, the computations by which expectations and sensory information are combined to refine perception remains an active area of investigation (de Lange et al., 2018, Press et al., 2020).

Bayesian inference describes the optimal means by which an observer can combine noisy sensory information with prior expectations to infer the state of the world. Strikingly, human behavior in perceptual tasks is often consistent with a Bayesian observer (e.g., Girshick et al., 2011; Jazayeri & Shadlen, 2010; Stocker & Simoncelli, 2006), engendering proposals that neural systems combine sensory inputs and expectations in this optimal fashion (for a recent review, see Aitchison & Lengyel, 2017).

Human neuroimaging has revealed that perceptual representations show characteristics of Bayesian inference. The result is a more precise representation that is biased towards prior expectations. In visual cortex, expected stimuli are more easily decoded from patterns of neural activity (Brandman & Peelen, 2017; Hindy et al., 2016; Kok et al., 2012) and reconstructed neural representations track prior expectations (Kok et al., 2013; van Bergen et al., 2015). However, prior studies have not independently manipulated sensory inputs and learned expectations in a quantitative manner, leaving unresolved the mechanism by which these cues are integrated.

We hypothesized that perceptual representations result from weighted averaging of feature estimates from sensation and expectation. In developing this hypothesis, we were inspired by the multisensory integration and cue combination literatures, which contain rigorous methods for evaluating fusion (Ban et al., 2012; Murphy et al., 2013). The key innovation of our study derives from examining fusion between an expectation cue and a sensory cue, whereas this prior work tested fusion between two sensory cues (e.g., depth from motion and disparity; Ban et al., 2012). After designing a learning paradigm that induces tone-based expectations about the sex of faces (Experiment 1), we tested for the fusion of these estimates of sex from tones and faces with model-based analyses of discriminability in behavior (Experiment 2) and the brain (Experiment 3).

## Methods

### Experiment 1

The purpose of Experiment 1 was to validate that we could establish a linear mapping between tones and faces through learning and that the resulting associations would induce expectations that bias behavior. Forty-eight human subjects participated in this study (28 female, mean age 19.6 years old). All had normal or corrected-to-normal vision. Informed consent was obtained according to a protocol approved by the Princeton University IRB.

Visual stimuli consisted of 41 sex-morphed face images (Zhao et al., 2011). These morphs were generated by interpolating features between a composite male face and a composite female face. The sex of the faces was coded using an arbitrary numerical index ranging from −1 to 1 in 0.05 increments, with −1 denoting the composite male face and 1 denoting the composite female face. Faces were presented centrally at fixation and spanned 4° of visual angle. We were not specifically interested in facial sex, but chose this domain because it is amenable to multivariate decoding from fMRI (Contreras et al., 2013; Kaul et al., 2011) and because face perception is linked with a well-defined cortical network (Dekowska et al., 2008).

Auditory stimuli consisted of 41 pure tones corresponding to musical notes D_1_ to B♭^7^ (36.7 to 3,951 Hz) in whole-step intervals. This tone space is perceptually uniform according to the MIDI pitch standard. The 41 tones were also assigned a numerical index ranging from −1 to +1 in 0.05 increments. For all experiments, the tone-face mapping was counterbalanced such that higher frequency tones were mapped to more masculine faces for half of the participants and to more feminine faces for the other half of participants. The amplitude of the tone stimuli was adjusted to correct for increasing subjective loudness with increasing pitch.

Participants completed 325 trials of a delayed estimation task (**Figure 1a**). On each trial, after being presented with a tone-face pair, participants had to morph a second face stimulus to match the sex of the face they had just seen as closely as possible. Participants morphed the face by dragging a mouse cursor to the left or right edge of the screen, which either smoothly incremented or decremented the sex of the face at the center of the screen. If a participant morphed the face to the end of the space, then morphing began to reverse direction. After identifying a desired face for their response, participants halted morphing by returning their cursor to the center of the screen and submitted their response by pressing the space bar.

**Figure 1.**
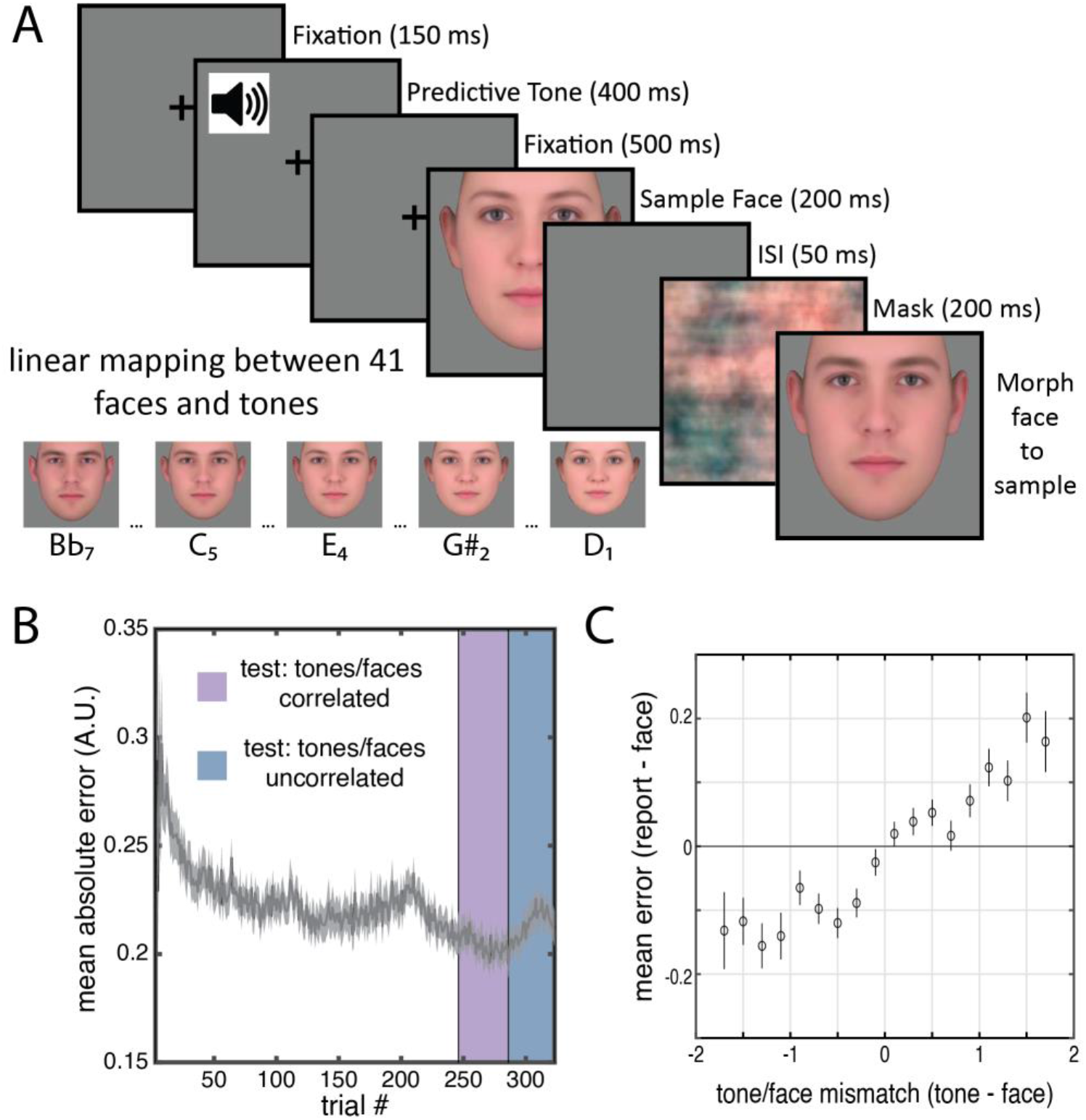
Learned tone-face associations bias behavioral reports. **(A)** Experiment 1 trial structure. On each trial, participants were presented with a pure tone and an image of a face drawn from a male-female continuum. At the end of the trial, participants continuously morphed a face through this space to match the sex of the face they had seen. Inset: example mappings for 5 tone face-pairs; in this example, lower notes were associated with more feminine faces. **(B)** Mean learning curve across participants. During an initial training phase (white region), the tones predicted the faces perfectly and participants received accuracy feedback. The congruent test phase (purple) was identical to this training phase, except that participants no longer received feedback. During the incongruent test phase (blue), the pairing of tone and face was random. Y-axis reflects error in units of facial sex: 0.05 units correspond to 1 step in the 41-step space. **(C)** Mean signed error (bias) as a function of tone-face mismatch during the incongruent test phase. Positive x-values indicate trials on which the tones predicted a more feminine face than was actually presented. Positive y-values indicate that participants reported a more feminine face than was actually presented. Both axes are differences in units of facial sex. Error bars are the standard error of the mean across participants.

The first 246 trials of this task constituted a training phase in which the tones were perfectly predictive of the faces and participants received feedback on their performance in the form of points. To encourage precision, points increased logarithmically as error approached zero, up to a maximum of 2,000. Negative points were awarded for errors greater than 0.30 units (6 steps in the 41-step space). Each tone-face pair was presented six times. Trial order was generated randomly for each participant.

Participants then completed two test phases (41 trials each) during which they no longer received feedback. In the first test phase, the tones remained perfectly predictive of the faces. In the second test phase, the mapping between tones and faces was randomly shuffled for each participant such that tone conveyed no information about the face. Each tone and face stimulus was presented once per test phase. Trial order was randomly generated for each participant.

Analysis of biases in participant’s reports during the training and first test phase revealed that a large proportion of the face stimulus space was approximately perceptually uniform. Across the interval from −0.7 to 0.7, mean absolute bias was 0.038 and the maximum absolute bias was 0.096, or less than one and two steps in the 41-step space, respectively. Stimuli were therefore restricted to this range in Experiments 2 and 3.

### Experiment 2

The purpose of Experiment 2 was to test whether tone and face information were integrated behaviorally in a manner consistent with fusion using a psychophysical task. Sixty new participants were recruited for this study (37 female, mean age 19.5). Participants were exposed to a linear mapping between tones and faces across 123 trials of a delayed estimation task identical to the training phase of Experiment 1 (**Figure 1a**), with three exposures of each tone-face pair. Trial order was generated randomly for each participant.

Participants then completed a discrimination task. On each trial, they were shown two tone-face pairs and asked to report whether the second face was more feminine than the first (**Figure 2a**). For the first pair, the tone continued to predict the sex of the face with 100% validity. The sex of this first tone-face pair was randomly assigned to 0.25, 0.20, 0.15, −0.15, 0.20, or 0.25 on each trial (‘g*’* in **Figure 2b**). For the second pair, however, the sex of the second tone and/or face was systematically manipulated in a manner that sometimes corrupted the predictive validity of the tone (**Figure 2b**): (1) On ΔFace trials, the second tone was identical to the first, but the second face differed in sex from the first by some increment. (2) On ΔTone trials, the second tone differed in sex from the first by some increment, but the second face was identical to the first. (3) On ΔCongruent trials, the second tone and face differed from the first tone and face by the same sex increment (the second tone on these trials was valid). (4) On ΔIncongruent trials, the second tone and face differed from the first tone and face by equal but opposite sex increments.

**Figure 2.**
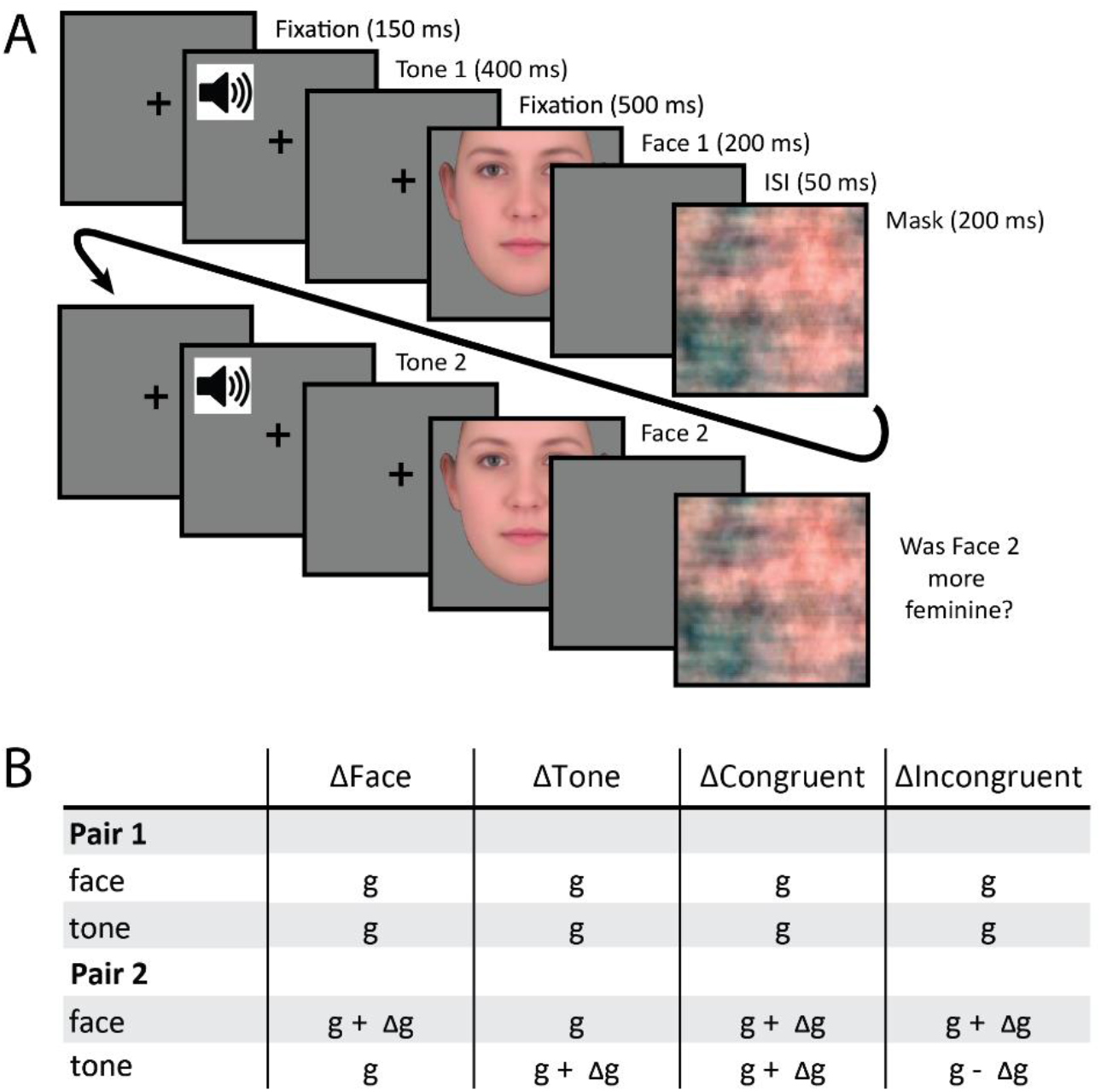
Discrimination task for testing fusion. **(A)** Discrimination task trial structure. On each trial, participants were presented with two tone-face pairs and reported whether the second face was more feminine than the first. **(B)** Discrimination task trial types. *g* refers to a point in the male-female space linked to a particular tone and face stimulus. Δ*g* is calculated separately for each condition and varies over trials according to a staircasing algorithm that identifies a 75% accuracy threshold (final Δ*g* = JND). For all trials, the first tone accurately predicted the first face. On ΔFace trials, the second face differed from the first. On ΔTone trials, the second tone differed from the first. On ΔCongruent trials, both the second tone and face differed from the first, and the second tone predicted the same sex as the second face. On ΔIncongruent trials, both the second tone and face differed from the first, but the second tone predicted a different sex than the face.

We measured the sensitivity of participants to increments in sex for each of these four trial types using separate staircases. Participants were not told about the existence of the different trial types or staircases. They began the discrimination task with 41 ΔCongruent trials. Sex increments on each trial were selected using a Bayesian adaptive algorithm (Watson & Pelli, 1983) to converge on the increment at which participants were correct 75% of the time. The purpose of this initial staircase was to avoid presenting invalid tones early on, which, coupled with the change in task phase, may have led participants to believe that the relationship between the tones and faces had changed. Results from this staircase were not analyzed. After the initial 41 trials, five additional and separate 41-trial staircases began concurrently. Depending on the trial type, the sex of the second tone and/or face were determined by increments from a ΔFace staircase, a ΔTone staircase, a ΔIncongruent staircase, or one of two ΔCongruent staircases. Two ΔCongruent staircases were included to help preserve the learned tone-face mapping by doubling the number of congruent trials. However, these two staircases were analyzed separately to equate statistical power across conditions. At the end of staircasing, the five estimated just noticeable differences (in units of sex space) were converted to sensitivity scores by taking their inverse (Ban et al., 2012).

By manipulating the predictive validity of the tones in this way, we were able to implement two tests for fusion. The first “quad-sum” test relates performance on ΔCongruent trials to performance on ΔFace and ΔTone trials. For a conservative null hypothesis, we still assume that participants use the tones when making their judgments, but that the sex conveyed by the faces and tones are encoded independently and are corrupted by independent sources of noise. Under these assumptions, the optimal solution is to recast the task as a discrimination problem in a space with two orthogonal cue axes (**Figure 3a)**. The discriminability of the two tone-face pairs on ΔCongruent trials is the hypotenuse (root quadratic sum) of the discriminability when only the tones or faces differ (**Figure 3b).** In the case of fusion, the sensory and expectation dimensions are not independent; observers take a weighted average of face and tone information for each pair to produce a single estimate of sex (**Figure 3c)**. Specifically, if the sex of the tone *t* is encoded with variance 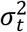 and the sex of the face *f* is encoded with variance 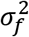, then the fused sex estimate on a particular trial will be drawn from a distribution with a mean

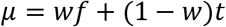

and reduced variance

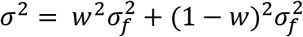

where *w* indicates the relative weighting of face and tone information. As a result of averaging, performance is suppressed in the ΔFace and ΔTone conditions because the difference along one dimension (i.e., face and tone, respectively) is diluted by averaging-in the other dimension that does not contain a difference (i.e., tone and face, respectively). Performance in the ΔCongruent condition should thus exceed the root quadratic sum of these suppressed levels (**Figure 3d**). Therefore, we computed the difference between the sensitivity in the congruent condition and the root quadratic sum of ΔFace and ΔTone sensitivity for each participant, and tested if the mean of this distribution was significantly greater than zero (two-tailed t-test):

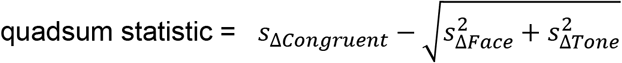

where *s* is sensitivity.

**Figure 3.**
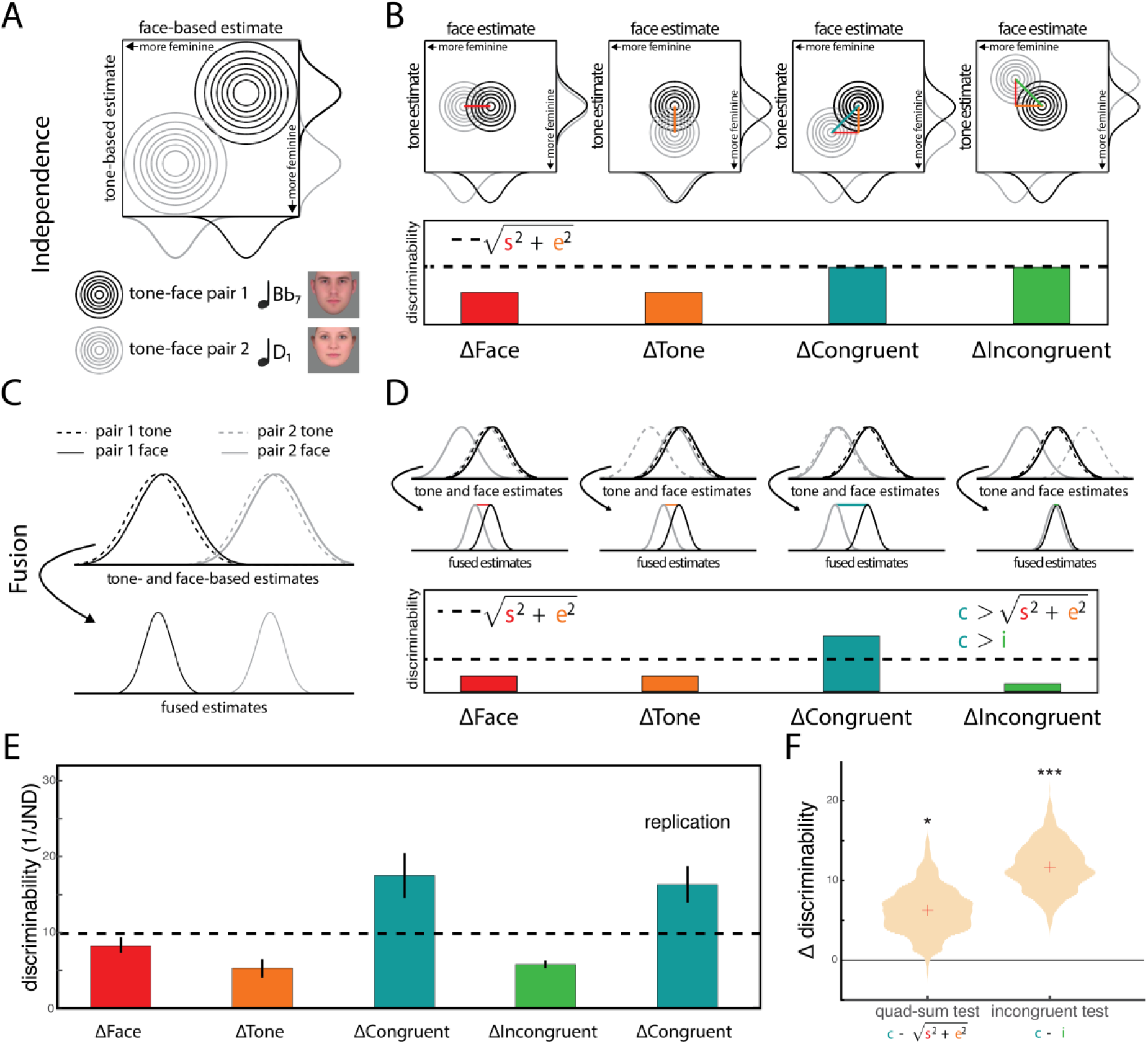
Theoretical predictions and behavioral fusion. **(A)** If the sex estimates elicited by the tones and faces are independent, the task can be recast as a linear discrimination problem in a 2D space. One axis represents sex estimates derived from the tones, and the other from the faces. Differences between the two pairs along either or both dimensions facilitate discrimination. The example depicts a ΔCongruent trial. **(B)** Predictions for each of the four trial types described in Figure 2b under independence. Because performance is proportional to the distance between means, performance in the ΔCongruent condition can be predicted by performance in the ΔFace and ΔTone conditions, according to the Pythagorean theorem (dotted line). Because cue axes are orthogonal, performance on ΔCongruent and ΔIncongruent trials is equal. **(C)** Under fusion, observers take a weighted average of sex estimates from the faces and tones, resulting in a 1D discrimination problem. **(D)** As a result, performance in the ΔFace and ΔTone conditions will be suppressed because an uninformative cue has been averaged-in (i.e., unchanging tone and face, respectively). Because both cues are informative, performance in the ΔCongruent condition is unaffected relative to independence, and thus exceeds the root quadratic sum (dotted line). ΔCongruent performance also exceeds the ΔIncongruent condition because the conflicting cues partially cancel each other out. **(E)** Sensitivity is measured as 1/JND, where JND reflects the Δ*g* in each condition that produced 75% discrimination accuracy. Error bars are the standard error of the mean across participants. **(F)** Mean of the two fusion metrics across participants (computed using the first of the two analyzed congruent staircases). Violin plots reflect the bootstrapped sampling distribution of the mean, in which participants were resampled with replacement to quantify reliability across participants. * p < 0.05, *** p < 0.001.

The second “incongruent” test for fusion compares performance on ΔCongruent vs. ΔIncongruent trials. An independence mechanism predicts that performance in the ΔCongruent and ΔIncongruent conditions should be equivalent because the distance between tone-face pairs in this bivariate space is the same (**Figure 3b).** In contrast, under fusion, the averaging of conflicting cues in the ΔIncongruent condition will reduce the differences between pairs and hamper discrimination relative to the ΔCongruent condition **(Figure 3d).** Therefore, we computed the difference in sensitivity between the ΔCongruent and ΔIncongruent conditions and tested if the mean of this distribution was significantly greater than zero (two-tailed t-test):

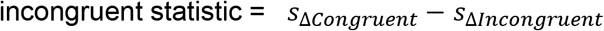

### Experiment 3

The purpose of Experiment 3 was to identify brain regions supporting neural fusion of tone and face information using multivariate fMRI. Thirty-two new participants were recruited for this study (20 female, mean age 21.8).The training task was identical to Experiments 1 and 2. Participants underwent training over the course of two days, completing 369 trials on the day prior to their scan and an additional 123 trials immediately before the scan.

In the scanner, participants were exposed to one tone-face pair per trial while performing an oddball cover task that demanded attention to the tone and face stimuli (**Figure 4a**). Oddball trials occurred ~18% of the time, containing either two tones or two faces in rapid succession in place of the typical one tone and one face. Participants were asked to report the presence of oddballs with a button press and these trials were discarded from further analysis. Participants completed eight fMRI runs of 98 trials each (18 oddball trials, 80 non-oddball).

**Figure 4.**
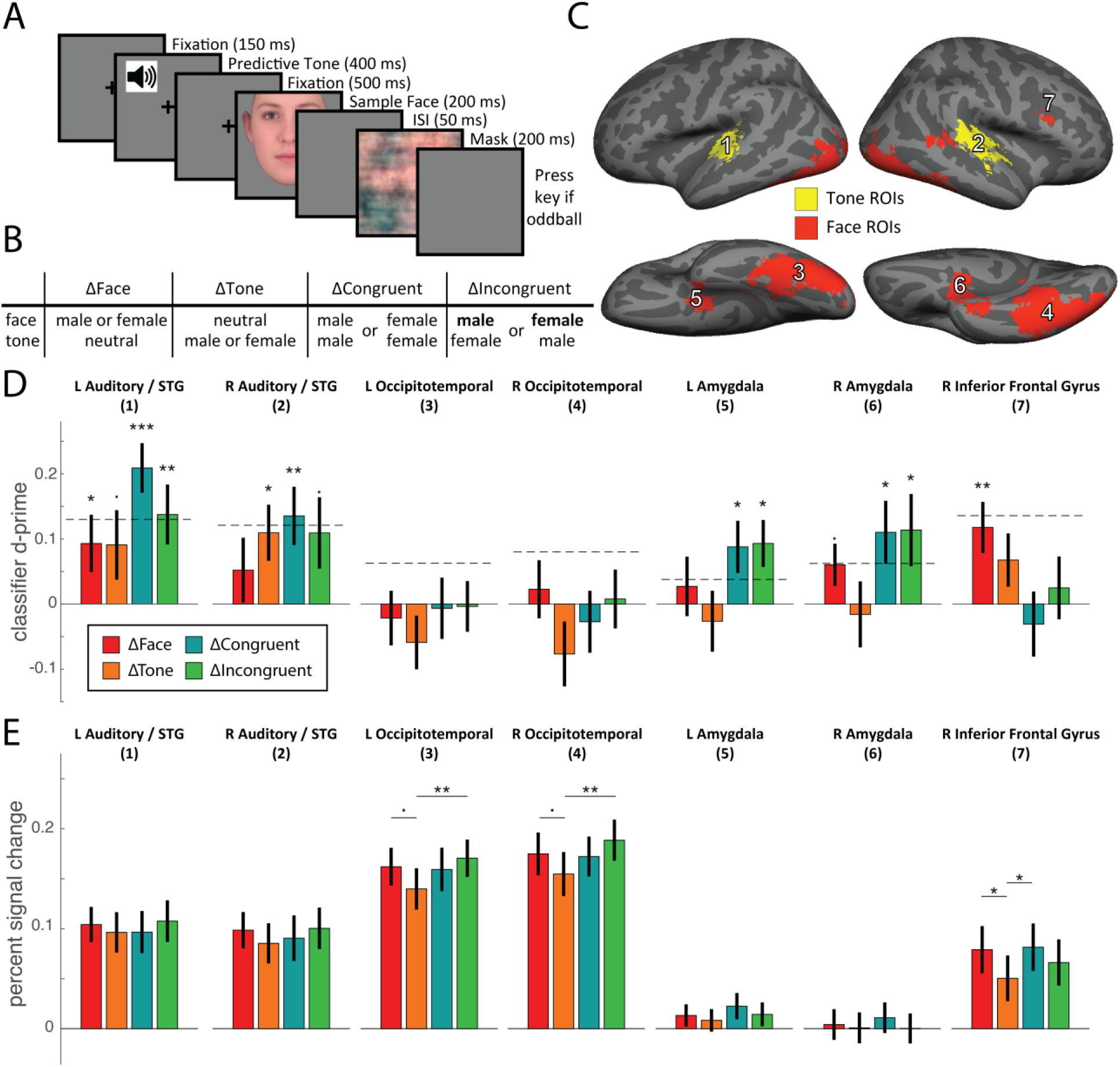
Neuroimaging design and results. **(A)** Each fMRI trial contained one tone-face pair. To ensure attention, participants pressed a key on infrequent “oddball” trials where either the tone or face was replaced by two rapid tones or faces, respectively, and otherwise withheld their response. Oddball trials were discarded from analysis. **(B)** There were four cue conditions: only the face indicating sex (ΔFace), only the tone indicating sex (ΔTone), the face and tone indicating the same sex (ΔCongruent), and the face and tone indicating different sexes (ΔIncongruent). Within each condition, the sex could either be male or female; ΔIncongruent trials were labeled based on the sex of the face. We assessed the neural evidence for facial sex in each condition by attempting to discriminate voxel patterns for male and female trials using multivariate pattern classification. **(C)** Regions of interest (ROIs) were generated using automated meta-analyses of published neuroimaging data (Yarkoni et al., 2011). (1) left auditory cortex / superior temporal gyrus (2) right auditory cortex / superior temporal gyrus (3) left occipitotemporal cortex (4) right occipitotemporal cortex (5) left amygdala (6) right amygdala (7) right inferior frontal gyrus. **(D)** Accuracy of the four classifiers (in units of d-prime) for each ROI. Dotted line indicates root quadratic sum of ΔFace and ΔTone d-prime, as in Figure 3. L/R indicate left/right hemisphere, STG = superior temporal gyrus. **(E)** Mean percent signal change for each condition and ROI, averaged across trials, voxels, and participants. Error bars are the standard error of the mean across participants. Lines and asterisks reflect paired t-tests. · p < 0.10, * p < 0.05, ** p < 0.01, *** p < 0.001.

The logic of Experiment 3 was similar to Experiment 2, but the discriminability of sex within each condition was examined differently: each trial contained one pair so we could estimate a neural representation of the sex of that tone-face combination from fMRI, and we calculated the discriminability of these representations across trials from the same condition. Specifically, non-oddball trials consisted of eight different trial types (10 trials each), resulting from the cross of sex (male or female) and condition (ΔFace, ΔTone, ΔCongruent, ΔIncongruent; **Figure 4b**). Across ΔFace trials the tone was neutral but the face conveyed sex; across ΔTone trials the tone conveyed sex but the face was neutral; across ΔCongruent trials both the tone and face conveyed sex and were consistent within trial; and across ΔIncongruent trials both the tone and face conveyed sex but were inconsistent within trial. Rather than fixing discriminability as we did in Experiment 2 (i.e., behavioral accuracy at 75%) and measuring the distance in stimulus space required, here we fixed the distance of the tones and faces in stimulus space and measured discrimination accuracy using multivariate pattern classifiers. Stimuli labeled “male” had a sex value of −0.6 (with +/− 0.1 units of jitter), stimuli labeled “female” had a sex value of 0.6 (with +/− 0.1 units of jitter), and neutral stimuli had a value of 0 (with +/ − 0.1 units of jitter).

Structural and functional MRI data were collected on a 3T Siemens Skyra scanner with a 16-channel head coil. Structural data were acquired using a T1-weighted magnetization prepared rapid acquisition gradient-echo (MPRAGE) sequence (1 mm isotropic). Functional data consisted of T2*-weighted multiband echo-planar imaging sequences with 48 oblique axial slices aligned to the AC-PC line acquired in an interleaved order (1,500 ms repetition time [TR], 40 ms echo time, 2 mm isotropic voxels, 96 × 96 matrix, 192 mm field of view, 64° flip angle). Data acquisition in each functional run began with 12 s of rest in order to approach steady-state magnetization. A B0 field map was collected at the end of the experiment.

The first four volumes of each functional run were discarded for T1 equilibration. Functional data were preprocessed and analyzed using FSL (www.fmrib.ox.ac.uk/fsl), including correction for head motion and slice-acquisition time, spatial smoothing (5-mm FWHM Gaussian kernel), and high-pass temporal filtering (128-s period). Data were manually inspected for motion artifacts, spikes, and low SNR.

We defined seven regions of interest (ROIs, **Figure 4c**), covering a broad swath of face- and tone-sensitive cortical areas, and tested whether their neural representations were consistent with fusion. ROIs were defined based on automated meta-analysis in Neurosynth (Yarkoni et al., 2011) using “face” and “tone” as search terms. ROIs were created by downloading statistical images from Neurosynth and binarizing the images such that significant voxels had a value of 1. Clusters with more than 100 voxels were saved as masks, registered to each participant’s functional space, and then re-binarized.

Classifier analyses were performed on the parameter estimates from a single trial GLM (Aly & Turk-Browne, 2016; Hindy et al., 2016), which contained 98 task-related regressors: one for every trial in the run, modeled as 1.5-s boxcars covering stimulus exposure. All regressors were convolved with a double-gamma hemodynamic response function. The six directions of head motion were also included as nuisance regressors. Autocorrelations in the time series were corrected with FILM prewhitening. Each run was modeled separately in first-level analyses. First-level parameter estimates were registered to the participant’s T1 image. For univariate analyses, parameter estimates were normalized to percent signal change by scaling with the min/max amplitude of the predicted effect, dividing by the run mean, and multiplying by 100 (http://mumford.fmripower.org/perchange_guide.pdf).

Classifier analyses were performed using custom scripts in MATLAB on individual runs and averaged across runs (Aly & Turk-Browne, 2016). For each participant, ROI, and condition (ΔFace, ΔTone, ΔCongruent, ΔIncongruent), we trained a regularized logistic regression classifier (penalty = 1) to distinguish voxel patterns of parameter estimates from “male” and “female” trials. Classifier performance was assessed using leave-one-out cross-validation (train on 19 trials, test on one). The average classifier accuracy across folds and runs was calculated separately for male and female test trials, and was converted to *d’* using the formula *z*(hit) – *z*(false alarm), where correct female test trials were coded as hits and incorrect male trials (i.e., labeled as female) were coded as false alarms. We then computed a neural version of the quadsum and incongruent fusion metrics for each participant and ROI by substituting classifier d-prime for sensitivity:

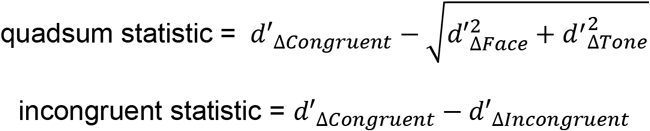

Finally, this fMRI design allowed us to perform an additional *a priori* “transfer” test for fusion (Ban et al., 2012; Murphy et al., 2013). Under fusion, estimates of sex from tones and/or faces are encoded in the same representational space. Therefore, a classifier trained to discriminate sex on ΔTone trials should be able to decode ΔFace trials, and a sex classifier trained on ΔFace trials should be able to decode ΔTone trials (**Figure 5c**). In contrast, when tone and face information are independent, a classifier trained on ΔFace trials should not successfully decode ΔTone trials, and vice versa. The transfer test statistic was thus the average d-prime of a classifier trained and tested in this manner. Specifically, within each run, classifiers were trained to decode male and female trials from the ΔFace condition and tested on the ΔTone condition, and vice versa. Generalization performance was averaged across these two folds and across runs.

**Figure 5.**
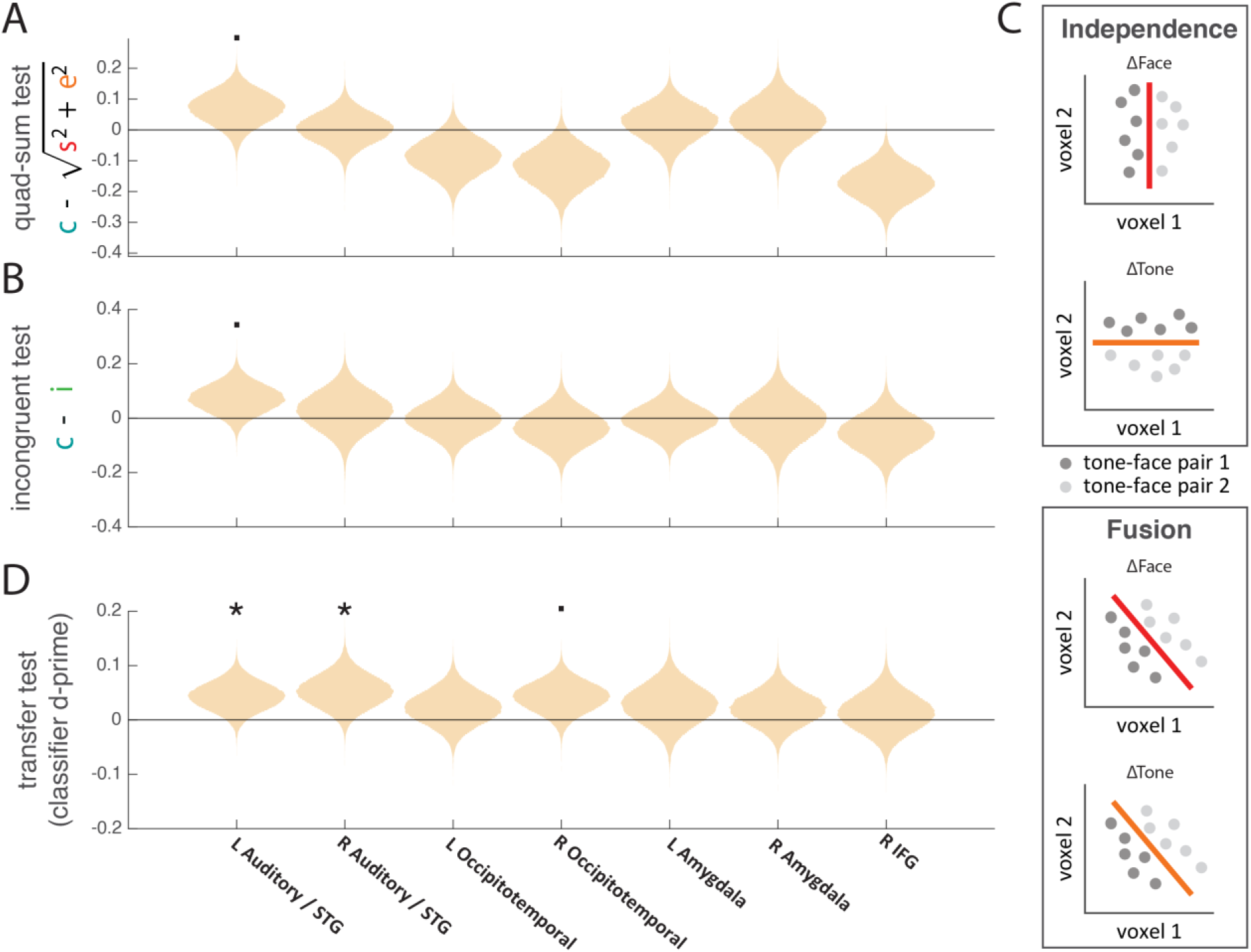
Classification-based neural fusion tests. **(A)** The mean quad-sum test statistic (ΔCongruent d-prime minus root sum-squared ΔFace and ΔTone d-prime) for each ROI. **(B)** The mean incongruent test statistic (ΔCongruent minus ΔIncongruent d-prime) for each ROI. **(C)** Rationale for the transfer test. Under independence, face and tone information are coded along orthogonal axis in state space. A classifier trained to discriminate male and female faces will fail to discriminate the corresponding tones (and vice versa). Under fusion, face and tone information are coded along a common axis, allowing a classifier to generalize from the ΔFace to the ΔTone condition (and vice versa). Dots reflect hypothetical trials in a 2-voxel state space; red and orange lines reflect trained classification boundaries. **(D)** The mean transfer test statistic for each ROI. Violin plots reflect the bootstrapped distribution of the mean. · p < 0.10, * p < 0.05.

The significance of each fusion metric for each ROI was assessed with random effects bootstrap resampling (Efron & Tibshirani, 1986), by sampling participants with replacement across 1000 iterations and calculating the proportion of iterations with a mean in the opposite direction as the true mean. The intuition behind this approach is that to the extent that an effect is reliable in the population, the participants should be interchangeable and a similar effect will be obtained regardless of which participants are sampled, resulting in a sampling distribution with low variance. To combine across fusion tests and control for multiple comparisons across ROIs, we also computed the probability that *any* of the 7 regions we investigated would display trending or significant effects for all three fusion tests by chance. To do this, we repeated the entire classification and bootstrapping procedure 1000 times, randomly permuting condition labels for each participant, and recorded the number of instances in which at least one ROI displayed p-values < 0.10 for all three fusion tests. This tested the null hypothesis that there was no meaningful pattern of classification performance across the four trial conditions.

We additionally used searchlight analyses to compute the three neural fusion metrics across cortex. The procedure was identical to that described above for the ROIs, except that parameter estimates were registered to 2-mm MNI space and analyses were repeated for all 27-voxel cubes (3 × 3 × 3) centered on voxels in cortex according to the Harvard-Oxford structural atlas (Desikan et al., 2006). Group analyses comparing each test to zero across participants were performed using random-effects nonparametric tests (as implemented by the ‘randomise’ function in FSL), corrected for multiple comparisons with threshold-free cluster enhancement (TFCE; Smith & Nichols, 2009).

We modeled the design and analysis of this imaging study after Experiment 2 because the independence model remains a strong null hypothesis. Indeed, any region that contains a mixture of face- and tone-selective voxels will display a pattern consistent with independence. The introduction of cue conflicts on ΔFace and ΔTone trials is a calculated design decision that allows us to test that superior performance in the ΔCongruent condition is, in fact, due to fusion, as described above.

## Results

### Experiment 1

We first established a novel set of expectations via associative learning in the domain of face perception. In the training phase, participants became more accurate across trials **(Figure 1b**, white region). The mean change in error from the first 20 trials to last 20 trials was 0.054, a significant decrease (*t*(47) = 5.019, *p* < 0.001).

In the test phase, we compared a congruent period in which the tones predicted the faces deterministically **(Figure 1b**, purple region) to an incongruent period in which there was no longer any relationship between tones and faces **(Figure 1b**, blue region). Error was significantly greater during the incongruent test than during the congruent test (*t*(47) = 2.824, *p* = 0.007). Errors during the incongruent test were influenced by the sign and magnitude of the tone-face mismatch. When the tone predicted a more feminine face than actually shown, participants tended to report a more feminine face, and vice versa (**Figure 1c**). The average within-subject correlation between tone-face mismatch and mean signed error across trials was *r* = 0.110, significantly greater than zero (*t*(47) = 10.204, *p* < 0.001).

Together, these results suggest that participants learned the mapping between tones and faces and that these associations were sufficient to generate expectations about facial sex that could bias behavior.

### Experiment 2

A new cohort of participants (N = 60) was exposed to a linear mapping between the tones and faces. We then tested whether expectations and sensory information were integrated in a manner consistent with fusion using psychophysical techniques originally developed for studying cue combination in depth perception (Ban et al., 2012; Murphy et al., 2013).

Consistent with a fusion mechanism: (1) sensitivity in both ΔCongruent conditions exceeded the root quadratic sum of ΔFace and ΔTone (quad-sum test, *t*(59) = 2.218, *p* = 0.030; *t*(59) = 2.053, *p* = 0.045), and (2) sensitivity in both ΔCongruent conditions exceeded ΔIncongruent (incongruent test, *t*(59) = 3.995, *p* < 0.001; *t*(59) = 4.359, *p* < 0.001, **Figure 3e-f**).

Together, these results provide evidence that expectations generated by recently learned cues are fused with sensory estimates.

### Experiment 3

We next tested for fusion of tones and faces in neural representations of sex. The pattern of classifier accuracy across conditions was most consistent with fusion in the left auditory cortex / superior temporal gyrus (STG) ROI, with a similar but weaker result in right auditory cortex / STG ROI (**Figure 4d**). The quad-sum (**Figure 5a**, *p* = 0.087) and incongruent (**Figure 5b**, *p* = 0.079) fusion tests trended toward significance in the left auditory cortex / STG ROI. The transfer fusion test (**Figure 5d**) was significant in left auditory cortex / STG (*p* = 0.042) and right auditory cortex / STG (*p* = 0.033).

Together, these analyses suggest auditory cortex / STG as a candidate region in which fusion may occur, particularly in the left hemisphere. Although the first two fusion tests were only marginally significant in this region (i.e., p < 0.10), the observation of a marginal or significant result for all three fusion tests in any of the seven ROIs was unlikely to have occurred by chance (*p* = 0.046, randomization test correcting for multiple comparisons).

The pattern of classification performance in left auditory cortex / STG was unrelated to the overall BOLD activity in each condition (**Figure 4e).** Indeed, repeated measures ANOVA revealed that percent signal change in this region was not modulated by condition (*F*(3,93) = 0.42, *p* = 0.740). Percent signal change was significantly modulated by condition in the inferior frontal gyrus (*F*(3,93) = 2.92, *p* = 0.038) and approached significance in left (*F*(3,93) = 2.41, *p* = 0.072) and right (*F*(3,93) = 2.61, *p* = 0.056) occipitotemporal cortex. Post-hoc t-tests revealed that this was due to lower percent signal change in the ΔTone condition (**Figure 4e)**.

Classification performance was poor in the two ventral face ROIs (all classifiers p > 0.05 vs. chance, **Figure 4d)**, perhaps due to weak topographic organizations for sex information. Classification in left and right amygdala was consistent with independence. In both regions, performance in the ΔCongruent and ΔIncongruent conditions was statistically indistinguishable (**Figure 5b)**, and while performance in the ΔCongruent was numerically greater than the ΔFace and ΔTone conditions, it did not exceed quadratic summation (**Figure 5a**). Right inferior temporal gyrus displayed an unexpected pattern in which performance in the ΔCongruent and ΔIncongruent conditions were statistically equivalent and tended to be worse than in the ΔFace and ΔTone conditions. The comparison of ΔCongruent vs ΔFace was significant (*p* = 0.018, randomization test, all other *p* > 0.10). Such a pattern could be generated by a region in which separate populations of voxels encode face and tone information and engage in mutually inhibitory interactions, though we are hesitant to interpret this one idiosyncratic finding.

To test for neural fusion outside of our ROIs, we ran an exploratory searchlight analysis across cortex for the three fusion tests. No regions passed the quad-sum or incongruent fusion tests after correcting for multiple comparisons. However, the transfer test searchlight revealed two large clusters in bilateral inferior temporal cortex, as well as smaller clusters in right auditory cortex (Heschl’s gyrus), and frontal, occipital, and parietal regions (**Figure 6, Table 1**). These results suggest that, after training, sensory stimuli and learned cues that evoke expectations about those stimuli can drive neural representations in a common manner throughout cortex.

**Figure 6.**
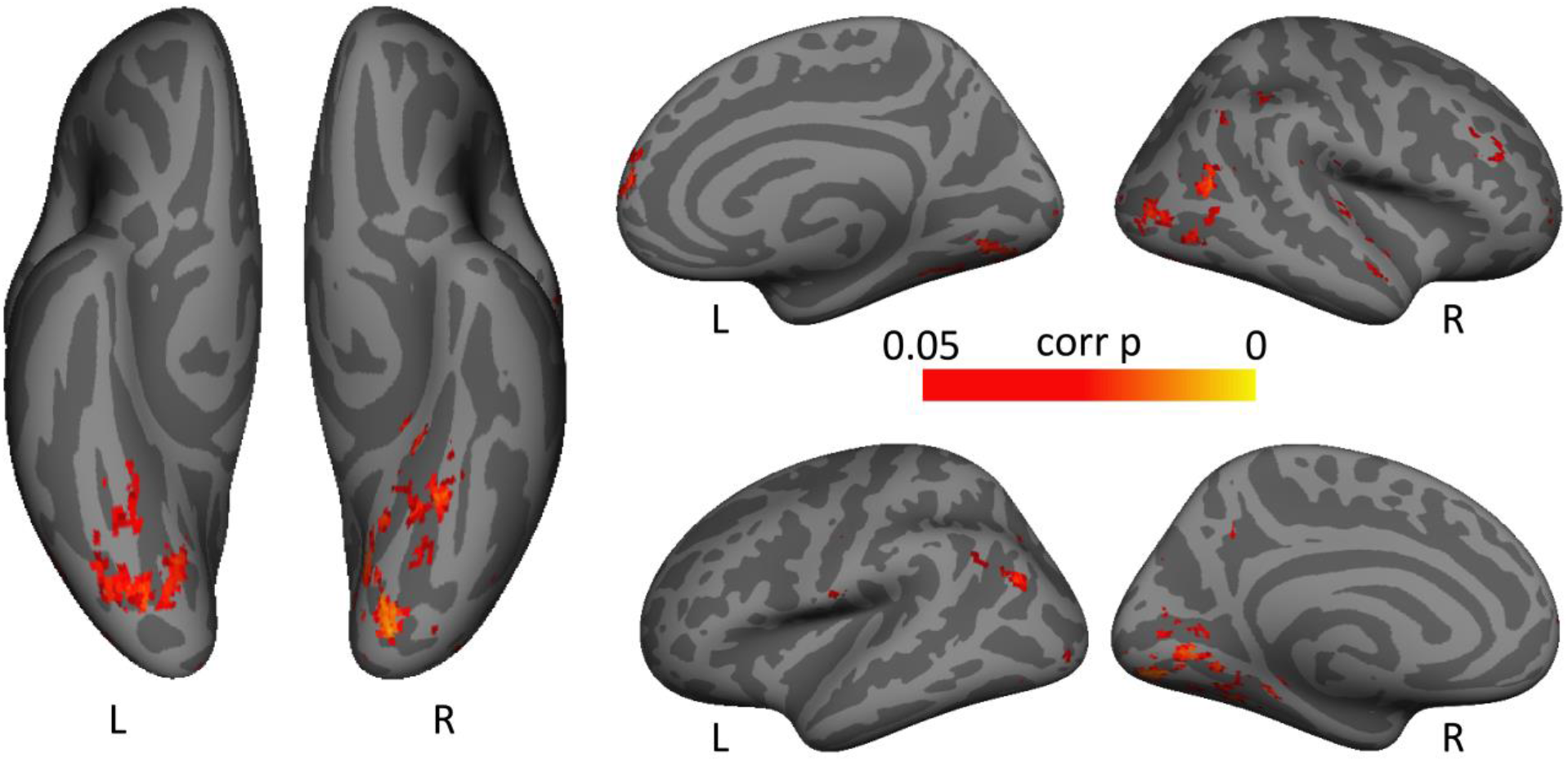
Results of transfer test searchlight. Thresholded statistical map of the transfer test searchlight analysis (corrected for multiple comparisons with TFCE). Patterns of activity surrounding voxels in the obtained clusters, many in visual cortex, shared a common code for sex from tones and faces. Positions of assigned values correspond to searchlight centers. L = left hemisphere, R = right hemisphere.

**Table 1.** Searchlight results from transfer test (clusters surviving correction)

| *Anatomical region* | *Hemi* | *Cluster size*<br>*(voxels)* | *Min p* | *MNI coordinates*<br>*(x y z)* |  |  |
| --- | --- | --- | --- | --- | --- | --- |
| Lingual Gyrus | L | 1349 | 0.011 | -14 | -88 | -8 |
| Occipital Fusiform Gyrus | R | 1053 | 0.012 | 34 | -76 | -4 |
| Paracingulate Gyrus | R | 134 | 0.025 | 10 | 54 | 14 |
| Lateral Occipital Cortex | L | 87 | 0.021 | -48 | -70 | 24 |
|  |  | 33 | 0.029 | -34 | -64 | 18 |
|  |  | 25 | 0.041 | -32 | -90 | -6 |
|  |  | 12 | 0.038 | -28 | -66 | 34 |
| Heschl's Gyrus | R | 70 | 0.032 | 54 | -18 | 14 |
| Precentral Gyrus | L | 34 | 0.040 | -62 | -4 | 12 |
| Middle Frontal Gyrus | R | 32 | 0.034 | 42 | 30 | 24 |
|  |  | 29 | 0.029 | 54 | 18 | 32 |
| Lateral Occipital Complex | R | 28 | 0.037 | 54 | -58 | 44 |
| Superior Temporal Gyrus | R | 27 | 0.040 | 52 | -6 | -16 |
| Occipital Pole | R | 22 | 0.040 | 20 | -90 | 6 |
| Central Opercular Cortex | L | 17 | 0.030 | -42 | -12 | 24 |
| Posterior Cingulate Gyrus | L | 14 | 0.029 | -6 | -36 | 12 |
|  |  | 11 | 0.041 | -10 | -50 | 32 |
| Frontal Pole | R | 12 | 0.041 | 30 | 50 | 6 |

## Discussion

This study provides evidence that observers incorporate expectations into perceptual processing by fusing them with sensory inputs. Conflicting sensory and expectation cues led to a specific pattern of behavioral deficits in perceptual decision-making, consistent with fusion models of cue integration in which feature estimates are averaged together. Pattern classifiers trained to perform an analogous set of discriminations based on neural activity displayed similar fusion in left auditory regions. These results provide evidence that fusion is instantiated at the neural level and suggests a computational mechanism by which expectations enhance the discriminability of perceptual representations (Brandman & Peelen, 2017; Hindy et al., 2016; Kok et al., 2012).

Note that while we did not observe a decrease in mean bold activity in auditory cortex, as observed in some other studies in regions which show enhanced discriminability of congruently cued stimuli (e.g. Kok et al., 2012), these results are not incompatible with proposals that expectations sharpen neural representations (de Lange et al., 2018; Kok et al., 2012). By analogy, highly successful models of visual attention marry mechanisms which can sharpen neural representations with inhibitory dynamics that maintain constant levels of overall neural activity (Reynolds & Heeger, 2009).

Previous work in multisensory integration and cue combination has demonstrated that humans can fuse highly stable cues that are genetically programmed or acquired over a lifetime of experience (Alais & Burr, 2004; Ban et al., 2012; Dekker et al., 2015; Ernst & Banks, 2002; Murphy et al., 2013; Nardini et al., 2010). Left auditory cortex including STG have been shown to be sensitive to the conjunction of familiar visual and auditory cues (e.g., video and audio of a person speaking, Callan et al., 2003; Hein et al., 2007; Kreifelts et al., 2007; Miller & D’Esposito, 2005). Here we show that the human brain flexibly leverages similar computational principles to integrate newly predictive information. This might explain how humans deploy recently learned environmental regularities in the service of faster and more accurate perceptual judgments (e.g., Esterman & Yantis, 2010; Turk-Browne, Scholl, Johnson, & Chun, 2010). Whether similar learning mechanisms govern both the rapid emergence of fusion in adults and the slower development of cue integration in children remains an intriguing and open question.

The present work has several limitations that should encourage further investigation. First, we explored fusion for only one type of feature: the sex of face stimuli. Future work could examine whether the present findings generalize to other features and feature-selective cortical areas. Second, we did not observe any evidence for fusion in ventral visual regions. This absence of an effect should be interpreted with caution because classification performance was generally poor in these regions. For example, the topography for identity-level facial information in these regions may be less clear than the tonotopic organization of auditory cortex. Alternative cover tasks that require more explicit judgements of face identity, as in previous work (Contreras et al., 2013; Kaul et al., 2011), may reveal clearer patterns of discrimination performance.

Bayesian inference provides a computational account of how expectations and sensory information interact in perception. The mechanism by which this integration is accomplished is likely to depend upon the type of expectation. For example, expectations may be embedded in the structural organization of cortex, or actively applied in the form of input from other brain regions (de Lange et al., 2018). Recent work suggests that expectations may be generated by the hippocampus when based on recently learned arbitrary associations (Hindy et al., 2016; Kok & Turk-Browne, 2018). This raises the possibility that the signatures of learned fusion in auditory cortex reported here may require hippocampal input.

## Author Contributions

M.F.P. and N.B.T.-B. designed experiments. M.F.P. collected and analyzed data. M.F.P. and N.B.T.-B. discussed the results and wrote the paper.

## Acknowledgments

This work was supported by a National Defense Science and Engineering Graduate Fellowship (M.F.P), US National Institutes of Health grant R01 EY021755 (N.B.T.-B.), and the Canadian Institute for Advanced Research (N.B.T.-B.). The authors thank Mariam Aly, Nick Hindy, Judy Fan, and Daniel Takahashi for helpful discussions.

